# The SecA motor generates mechanical force during protein translocation

**DOI:** 10.1101/2020.04.29.066415

**Authors:** Riti Gupta, Dmitri Toptygin, Christian M. Kaiser

**Affiliations:** CMDB Graduate Program, Johns Hopkins University; Department of Biology, Johns Hopkins University; Department of Biophysics, Johns Hopkins University

## Abstract

The Sec translocon moves proteins across lipid bilayers in all cells. The Sec channel enables passage of unfolded proteins through the bacterial plasma membrane, driven by the cytosolic ATPase SecA. Whether SecA generates mechanical force to overcome barriers to translocation posed by structured substrate proteins is unknown. Monitoring translocation of a folded substrate protein with tunable stability at high time resolution allowed us to kinetically dissect Sec-dependent translocation. We find that substrate unfolding constitutes the rate-limiting step during translocation. Using single-molecule force spectroscopy, we have also defined the response of the protein to mechanical force. Relating the kinetic and force measurements revealed that SecA generates at least 10 piconewtons of mechanical force to actively unfold translocating proteins, comparable to cellular unfoldases. Combining biochemical and single-molecule measurements has thus allowed us to define how the SecA motor ensures efficient and robust export of proteins that contain stable structure.

## Introduction

The universally conserved Sec translocon^1^ transports proteins across membranes in all cells. In bacteria, the SecYEG complex forms a protein conducting channel in the bacterial plasma membrane^2^. While integral inner membrane proteins pass through this channel as they are being synthesized by the ribosome, a large fraction of secreted and outer membrane proteins are exported after their synthesis is complete. This post-translational translocation process is driven primarily by the translocon-associated ATPase, SecA^2–6^.

During translocation, SecA undergoes repeated cycles of ATP binding, hydrolysis, and ADP/Pi release. How the chemical energy from ATP hydrolysis is converted into mechanical work that drives translocation is not well understood. “Power stroke” models posit that a structural element in SecA, termed the two-helix finger, inserts into the SecY channel upon ATP binding, dragging the translocating polypeptide along with it^7^,^8^. Two other SecA domains, the polypeptide crosslinking domain and the nucleotide binding domain 2, form a clamp that subsequently closes around the substrate, preventing backsliding as the two-helix finger resets after ATP hydrolysis^9^. In an alternative “Brownian ratchet” model, bulky residues on the cis side of the translocon trigger ATP binding to SecA, which then elicits conformational opening of the SecY channel that permits passive diffusion of the translocating polypeptide^10^. ATP hydrolysis results in channel closure, rectifying the progress of translocation that has occurred spontaneously. Direct experimental evidence for either mechanism remains sparse.

Tertiary structure in Sec substrate proteins interferes with translocation because the central aqueous channel in the SecYEG pore is narrow^11,12^, permitting only unfolded polypeptides to pass through^13^. Cellular proteins destined for export through the Sec translocon are thought to be kept in a largely unfolded state with the help of molecular chaperones^13,14^ However, even in the presence of chaperones that serve to prevent premature folding in the cytosol^15^, substrate proteins can acquire stable structure. For instance, the precursor outer membrane protein A (pOA), a Sec substrate, exhibits substantial secondary and tertiary structure in the presence of the chaperone SecB^16^. Such structures must be unfolded prior to translocation, presumably in an active process driven by ATP hydrolysis^14^ It has been suggested that this unfolding is accomplished through mechanical force generated by the SecA motor^7,9,14^

Mechanical force acts as a denaturant that destabilizes folded proteins. Cellular machinery has been demonstrated to utilize mechanical force for protein unfolding^17,18^ and disaggregation^19^. The translocon motor SecA has similarly been suggested to convert chemical energy from ATP hydrolysis into mechanical work that might help to unravel folded substrate proteins^6,7,20,21^ but the magnitude of the relevant forces remains unknown. Conflicting experimental findings argue for a regulatory role of ATPase activity^10^. As such, it remains unclear whether SecA is capable of generating mechanical forces, and whether such forces would be large enough to substantially accelerate substrate protein unfolding.

Here, we combined biochemical translocation assays and single-molecule force spectroscopy experiments to determine whether the SecA translocation motor can act as a mechanical unfoldase. A substrate protein whose mechanical stability can be tuned with small-molecule ligands provided a defined roadblock that must be unfolded prior to passing through the SecYEG channel. Continuous real-time translocation measurements, analyzed with a novel kinetic model, allowed us to quantify the unfolding kinetics of during translocation. Using optical tweezers experiments, we quantitatively defined the unfolding rate of the roadblock as a function of mechanical load. By combining these analyses, we estimate the force exerted on the substrate protein during translocation. Collectively, our analyses suggest that SecA acts as a power-stroke motor that generates mechanical force to aid the unfolding of substrate proteins, ensuring efficient export of proteins containing stable tertiary structure.

## Results

### Real-time measurements enable kinetic dissection of translocation

Stably folded structures of Sec-substrates must be unfolded prior to translocation, slowing the progress of the reaction^14,22^. To obtain a quantitative understanding of substrate unfolding at the translocon, we conducted translocation experiments with dihydrofolate reductase (DHFR). DHFR has been widely used to study mitochondrial protein import^23–26^ and Sec-dependent translocation^7,27^. For our studies, we utilized the murine enzyme (mDHFR, Figure 1A). The natural substrates of the enzyme, dihydrofolate and nicotinamide adenine dinucleotide phosphate (NADPH), and the inhibitor methotrexate (MTX) have been shown to increase the thermodynamic stability of mDHFR^24,28,29^. The protein thus functions as a translocation roadblock with tunable stability.

**Figure 1.**
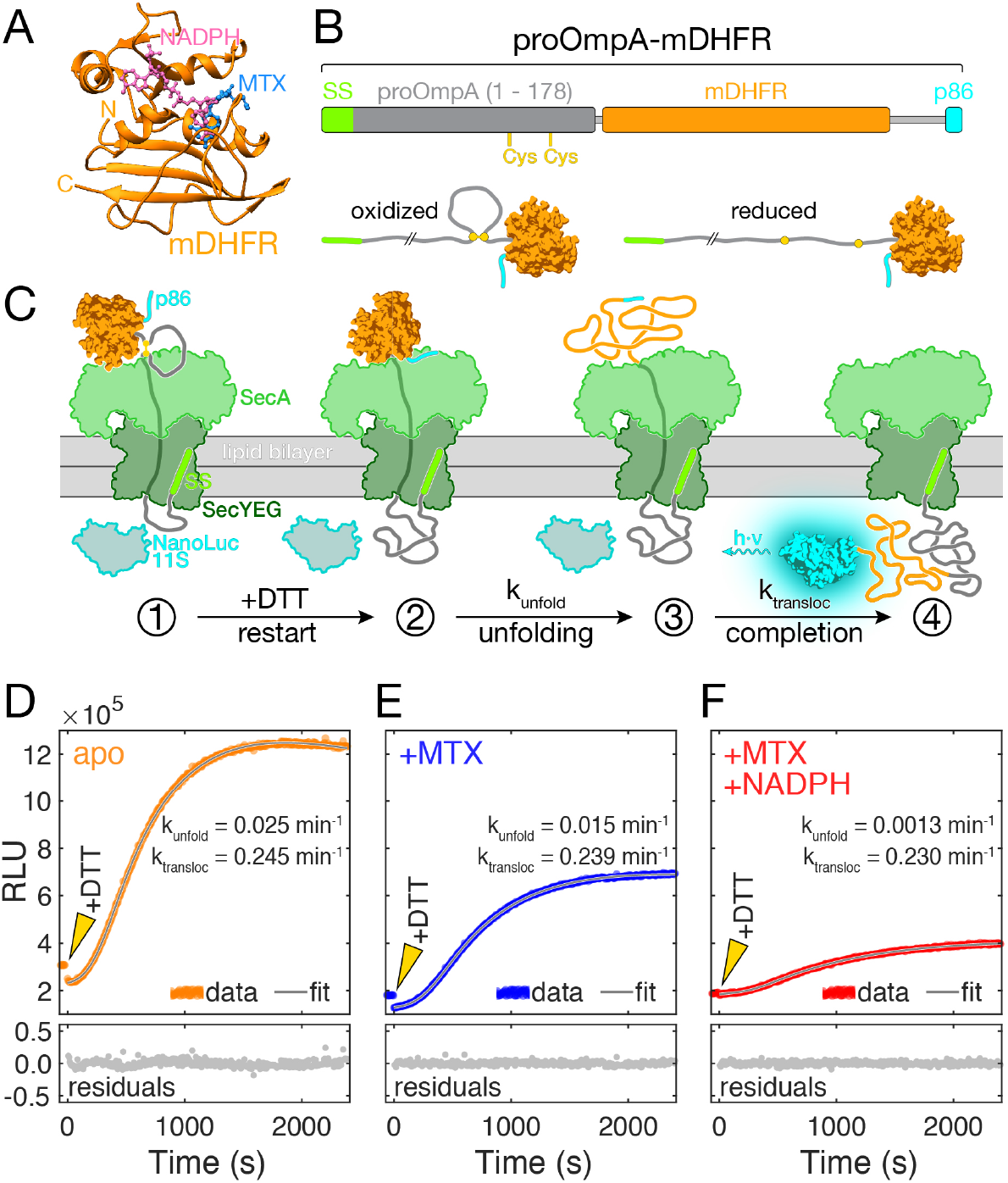
Synchronized real-time measurements enable kinetic dissection of translocation and unfolding. **A.** Crystal structure of mDHFR bound to the ligands MTX and NADPH (pdb: 1u70). **B.** Schematic of the chimeric proOmpA-mDHFR translocation substrate (top). Two cysteine residues engineered in the proOmpA N-terminal fragment allow the reversible formation of a looped structure (bottom). **C.** Schematic of the translocation experiment. Initial translocation of the oxidized proOmpA-mDHFR substrate results in complexes stalled at the disulfide loop (➀). The NanoLuc 11S protein (cyan/grey) is encapsulated inside the proteoliposomes and separated from the complementing p86 peptide (cyan) by the lipid bilayer. Upon DTT addition, translocation proceeds until mDHFR reaches the translocon (➁). After unfolding (➂), translocation is completed, resulting in reconstitution of luciferase activity and light emission (➃). **D – F.** Representative examples of real-time translocation recordings for apo, MTX and MTX+NADPH conditions, showing the recorded data (relative light units, RLU) as colored circles and the fit to the kinetic model (see main text for details) as a grey line. Residuals (bottom) indicate that the fit describes the data well. Addition of DTT at t = 0 (arrowheads) restarts translocation. Ligands decelerate unfolding of mDHFR (k_unfold_), resulting in reduced rates and amplitudes of the translocation signal. The translocation rates (k_transloc_) are similar for all conditions.

We constructed a chimeric translocation substrate protein by fusing mDHFR to the N-terminal 178 amino acids of pOA that included the signal sequence (Figure 1B). Engineered cysteine residues at positions 122 and 152 in pOA allowed us to form an intramolecular loop by oxidation (Figure 1B, Supplementary Figure 1). The disulfide loop creates a physical barrier to translocation progress^7^,^30^, stalling the reaction at a position close to residue 122. When a reducing agent such as dithiothreitol (DTT) is added, translocation resumes and the C-terminal portion of the fusion protein, including mDHFR, is imported (Supplementary Figure 1).

To analyze translocation kinetics at high time resolution, we utilized a recently described assay^31^ that continuously follows the reaction progress in real-time through light emission by the NanoLuc luciferase. NanoLuc can be split into two asymmetric fragments p86 (11 amino acids) and 11S (159 amino acids), neither of which has significant enzymatic activity. The p86 peptide tag binds 11S with very high affinity (K_D_ = 700 pM)^32^, restoring luciferase activity. We genetically fused the p86 peptide to the C-terminus of pOA-mDHFR (Figure 1B) and encapsulated 11S inside proteoliposomes containing reconstituted SecYEG and SecA (Figure 1C).

When the oxidized pOA-mDHFR fusion protein engages with the translocon complex, the pOA portion is translocated up to position 122, where the disulfide loop blocks further translocation (Figure 1C, ➀). Translocation resumes after the addition of reducing agent until the folded mDHFR reaches the translocon (Figure 1C, ➁). After mDHFR unfolding (Figure 1C, ➂), translocation is completed, and luciferase activity is restored (Figure 1C, ➃), reporting on translocation progress.

Figure 1D shows a representative example of a real-time translocation measurement of pOA-mDHFR at room temperature in the absence of stabilizing ligands (“apo”). After addition of DTT to stalled translocon substrate complexes (arrowhead in Figure 1D), luciferase activity began to increase after an initial delay and leveled off within approximately 30 minutes. To extract kinetic information from these measurements, we developed a detailed model (Materials and Methods, and Supplementary Information). The model takes into account the rates for mDHFR unfolding and translocation (k_unfold_, k_transioc_), as well as luciferase substrate depletion and an additional rate that accounts for loss of translocatable substrate over time (“incapacitation”, see next section, Supplementary Figure S2, and Supplementary Information). The model fits the data well (Figure 1D, and Supplementary Figure S3), indicating that it provides a suitable description of our experimental system.

We determined the rates of unfolding and translocation to be k_unfold_(apo) = 0.025 min^−1^ and k_transloc_(apo) = 0.245 min^−1^ (see Table 1 and Supplementary Table S1 for summaries of the fit parameters and their standard deviations). Previously reported translocation rates measured at 30°C using similarly reconstituted translocon complexes ranged from 0.1 to 1.2 min^−1^ for substrate proteins that are not expected to contain stable structures^33^. The translocation rate that we extract from our kinetic data is consistent with these published values that were determined in traditional protease protection assays.

The results from our real-time translocation measurements indicate that unfolding of apo-mDHFR takes approximately 10 times longer than the actual translocation reaction. This result suggests that mDHFR is stably folded and poses a significant barrier to translocation, even in the absence of ligands. After mDHFR unfolding, the protein is fully translocated into the interior of the proteoliposomes. Following translocation continuously (sampling at a rate of ~0.2 Hz in our measurements) makes it possible to dissect the kinetic components of the process.

**Table 1.** Summary of fit parameters from analyzing real-time translocation measurements. The table shows the mean ± standard deviation of the unfolding, translocation and incapacitation rates (k_unfold_, k_transloc_, k_incap_) for three experimental conditions.

| Condition | $k_{\text{unfold}} (\text{min}^{-1})$ | $k_{\text{transloc}} (\text{min}^{-1})$ | $k_{\text{incap}} (\text{min}^{-1})$ |
| --- | --- | --- | --- |
| apo | $0.0251 \pm 0.000154$ | $0.245 \pm 0.000589$ | $0.0658 \pm 0.000107$ |
| MTX | $0.0115 \pm 0.000114$ | $0.239 \pm 0.000826$ | $0.0729 \pm 0.000187$ |
| M+N | $0.00130 \pm 0.000025$ | $0.230 \pm 0.00193$ | $0.0547 \pm 0.000411$ |

### Structure stabilization slows translocation

A previous study, using protease protection to follow Sec-dependent import of a DHFR-fused substrate protein, indicated that the translocation of mDHFR is not significantly impaired by the ligand MTX alone, whereas a combination of MTX and NADPH slowed down the reaction^27^. In the presence of MTX, we observed a reduced overall rate of protein import into proteoliposomes (Figure 1E). Analyzing the time trajectories obtained with MTX-bound pOA-mDHFR, we find that the unfolding rate is reduced to k_unfold_(MTX) = 0.012 min^−1^, approximately half the value obtained for the apo-protein. We therefore observe a small but measurable deceleration of unfolding in the presence of the ligand.

A more pronounced effect on the overall rate of substrate import is observed when both MTX and NAPDH (M+N) are added. The unfolding rate is drastically reduced in this case to k_unfold_(M+N) = 0.0013 min^−1^, almost 20-fold lower than the rate for apo-mDHFR unfolding (Figure 1F). The finding that simultaneous binding of MTX and NADPH stabilizes native mDHFR against translocon-mediated unfolding is consistent with previously reported results^27^. However, the time resolution afforded by our real-time translocation measurements allows us to separate unfolding from translocation kinetically and obtain rates for the two processes individually.

Unfolding of mDHFR results in ligand dissociation^34^ Therefore, the translocated polypeptides are chemically identical in liganded and ligand-free conditions, avoiding the convoluting effects of polypeptide sequence on translocation rates^33,35^. Based on these considerations, the rates of translocation after unfolding of mDHFR is expected to be constant in all of our measurements. Indeed, fitting our model to the data yields translocation rates that are very similar in the presence (k_transloc_(MTX) = 0.239 min^−1^ and k_transloc_(M+N) = 0.230 min^−1^) and absence of ligands (k_transloc_(apo) = 0.249 min^−1^). The agreement of translocation rates determined in our assay validates the model that we use to interpret our translocation measurements.

While the translocation rates are similar, we noted that the amplitudes of the observed signal progressively decrease from the apo to the MTX and the M+N condition. The decrease in the final luminescence signal suggests that unfolding competes with a process that renders the system incapable of translocation. We therefore termed this process “incapacitation”. Incomplete translocation, indicating substrate attrition, is commonly observed in translocation measurements (see ref.^33^, and references therein), but typically not accounted for. The incapacitation rates (k_incap_) are similar for all experimental conditions (k_incap_(apo) = 0.066 min^−1^, k_incap_(MTX) = 0.073 min^−1^, k_incap_(M+N) = 0.055 min^−1^; see Table 1 for standard deviations). However, because ligand binding decreases the unfolding rate while the rate of irreversible incapacitation remains constant, reduced amounts of translocated product are observed.

Taken together, we find that the ligands MTX and NADPH stabilize mDHFR against unfolding by the translocon. The experimental design employed here, together with the detailed kinetic model that we have developed, allows us to quantify the unfolding rate at the translocon, which decreases in response to ligand binding, while the rate of translocation remains constant. Unfolding by the translocon has been suggested to be aided by mechanical force generated by SecA using ATP as fuel^7,14^ but quantitative measurements supporting this idea are lacking. Determining the unfolding kinetics of mDHFR under mechanical load would permit a quantitative comparison with the unfolding rates observed during translocation to test this hypothesis.

### mDHFR is mechanically stable

To determine the mechanical stability of mDHFR, we characterized its unfolding by singlemolecule force spectroscopy with optical tweezers, a powerful tool for characterizing folding energy landscapes^36–38^. To make the protein amenable to mechanical manipulation, we genetically engineered sites for the attachment of molecular handles^39^ that link the termini of the protein to two beads, one held in a micropipette and the other in an optical trap (Figure 2A). By moving the optical trap away from the micropipette, the tethered protein is subjected to mechanical force that acts as a denaturant, biasing the molecule toward unfolding.

**Figure 2.**
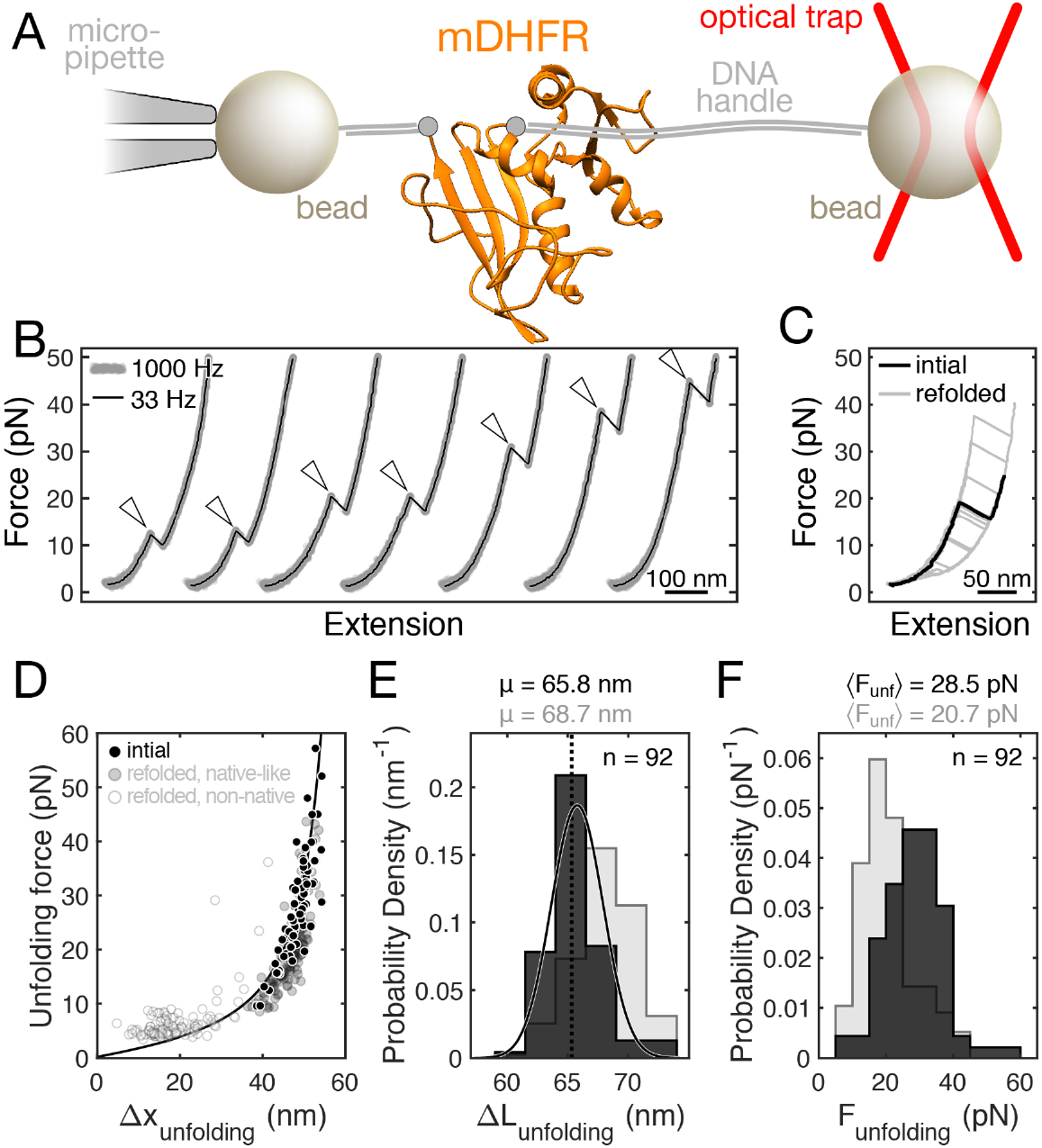
apo-mDHFR is mechanically stable and structurally heterogeneous. **A.** Cartoon schematic of optical tweezers experiment (not to scale). mDHFR is tethered by its termini between two beads for mechanical manipulation with an optical tweezers. **B.** Example force-extension curves showing initial unfolding events (arrowheads) for seven apo-mDHFR molecules, plotted with an offset along the horizontal axis for clarity, and sorted by unfolding force in ascending order. **C.** Force-extension curves for the initial (black line) and subsequent (grey lines) unfolding of a single apo-mDHFR molecule. Some of the curves recorded after refolding resemble the initial one, others occur at low force with shorter extension changes. **D.** Scatter plot of initial (black circles) and subsequent (open and filled grey circles) unfolding events for 92 apo-mDHFR molecules. Each dot represents an unfolding event, characterized by the unfolding force (y-axis) and the extension change upon unfolding (Δx_unfolding_, x-axis). Subsequent unfolding events are colored depending on whether they exhibit native-like (filled grey circles) or non-native (open grey circles) characteristics. Black line: worm-like chain model calculated for native mDHFR unfolding. **E, F.** Histograms showing the distributions of contour length changes (E) and unfolding forces (F) for initial (dark grey) and native-like (light grey) unfolding transitions. A fit of the contour length changes from initial events to a normal distribution (solid line) yields a mean (μ) close to the expected value (dotted line) for mDHFR unfolding. The unfolding forces from initial events exhibit a distribution with a tail at high force that is uncharacteristic for unfolding from a single state. The respective distributions for native-like transitions overlap with those for initial events, but are statistically distinct, indicating inefficient refolding to the initial structure. Average unfolding forces (〈F_unf_〉) are indicated on top of panel F.

We carried out “force ramp” experiments by moving the optical trap at a constant velocity of 150 nm/s, while recording the force and the molecular extension. In the resulting force-extension curves, mDHFR unfolding manifests as a “rip” (arrowheads in Figure 2B), a sudden increase in molecular extension. Figure 2B displays representative examples of initial unfolding events for seven mDHFR molecules. Unfolding occurs in a broad range of forces from approximately 10 to 55 piconewtons (pN). The mechanical stability of apo-mDHFR is similar to that of several other globular proteins that have been subjected to similar measurements, such as ribonuclease H^40^, calmodulin^41^, T4 lysozyme^39^, or elongation factor G^42^, which unfold within this force range. Interestingly, a previous study using atomic force microscopy (AFM) indicated that Chinese hamster DHFR is mechanically weak^43^. The difference in mechanical behavior between the mouse and hamster proteins might be due to structural differences that have been observed by crystallography for closely related DHFR orthologs^44^

Most unfolding transitions observed in our experiments with apo-mDHFR occur in one apparent step. However, in some events occurring at low forces, a transient unfolding intermediate is observed (Supplementary Figure S4A). The estimated contour length change for the transition from the native to the intermediate state is approximately 20 to 25 nm (Supplementary Figure S4B), similar to an intermediate detected in AFM experiments of mDHFR^45^. The total length change for the initial unfolding events, calculated using a worm-like chain model^46^ matched the expectation for mDHFR unfolding (Figure 2E, black dots and black line). The total unfolding length changes (ΔL_unfolding_) are distributed around ΔL_unfolding_ = 65.8 nm (Figure 2C, black histogram), very close to the expected value of ΔL_unfolding_(calc) = 65.4 nm (Figure 2C, dotted line) calculated from the mDHFR crystal structure^44^ Taken together, the observed unfolding events are consistent with complete unfolding of natively structured mDHFR.

After the initial unfolding event, we relaxed the force to allow refolding. Stretching the protein after holding it at a low force of 2 pN for 15 seconds yielded force extension curves (Figure 2C) with heterogeneous transitions that were either distinct from the initial ones or had similar properties (Figure 2D, white and grey dots, respectively). Even the events that showed a length change similar to the initial unfolding (Figure 2D, grey dots) exhibited contour length and unfolding force distributions that overlapped with, but were distinct from, the initial events (Figure 2E and F). We conclude that mDHFR does not refold efficiently under our experimental conditions and therefore focused our analysis on the initial unfolding events.

Unfolding force distributions obtained in force ramp experiments contain information about the underlying molecular process. For an unfolding process with one rate-limiting step, the continuously increasing unfolding rate results in a characteristic skewed distribution of the unfolding force (Supplementary Figure S5A). The unfolding force distribution of apo-mDHFR does not exhibit this characteristic shape (Figure 2F). Instead, the distribution is consistent with at least two distinct barriers, resulting either from an equilibrium of states with distinct stabilities or from alternative unfolding pathways (Supplementary Figure S5). Given the distinct distributions observed for initial and subsequent events (Figure 2F), our results likely reflect the population of several native states in mDHFR that equilibrate slowly, as has been observed for the orthologous *E. coli* DHFR^47^ This structural heterogeneity makes it challenging to extract kinetic parameters that govern unfolding.

Taken together, our single-molecule unfolding experiments suggest that native mDHFR exists in a conformational equilibrium, populating at least two states that differ in their mechanical stabilities. Either state exhibits significant mechanical stability, which explains the observed slow unfolding during translocation. However, the apparent structural heterogeneity of apo-mDHFR hampers a quantitative analysis of its mechanical properties.

### Ligands increase mDHFR mechanical stability

The effects of ligand binding on the mechanical stability of mammalian DHFR are not well understood. Conflicting results from atomic force microscopy (AFM) experiments suggest either stabilization of the native state^43^, or no effect on native state stability but stabilization of an unfolding intermediate^45^. Force ramp experiments in the presence of 10 μM MTX (Figure 3A) yielded transitions with contour length changes of ΔL_unfolding_ = 63.5 nm, close to the expected value (Figure 3B). As observed with apo-mDHFR, some unfolding traces recorded in the presence of MTX exhibit a transient intermediate (Supplementary Figure S4). However, the unfolding forces in the presence of MTX are higher than those of apo-mDHFR, ranging from 33 to 58 pN (Figure 3C). Compared to the scenario of apo-mDHFR, the shape of the unfolding force distribution for MTX-bound mDHFR more closely matches the expectation for unfolding from a single well-defined state (see Supplementary Figure S5).

**Figure 3.**
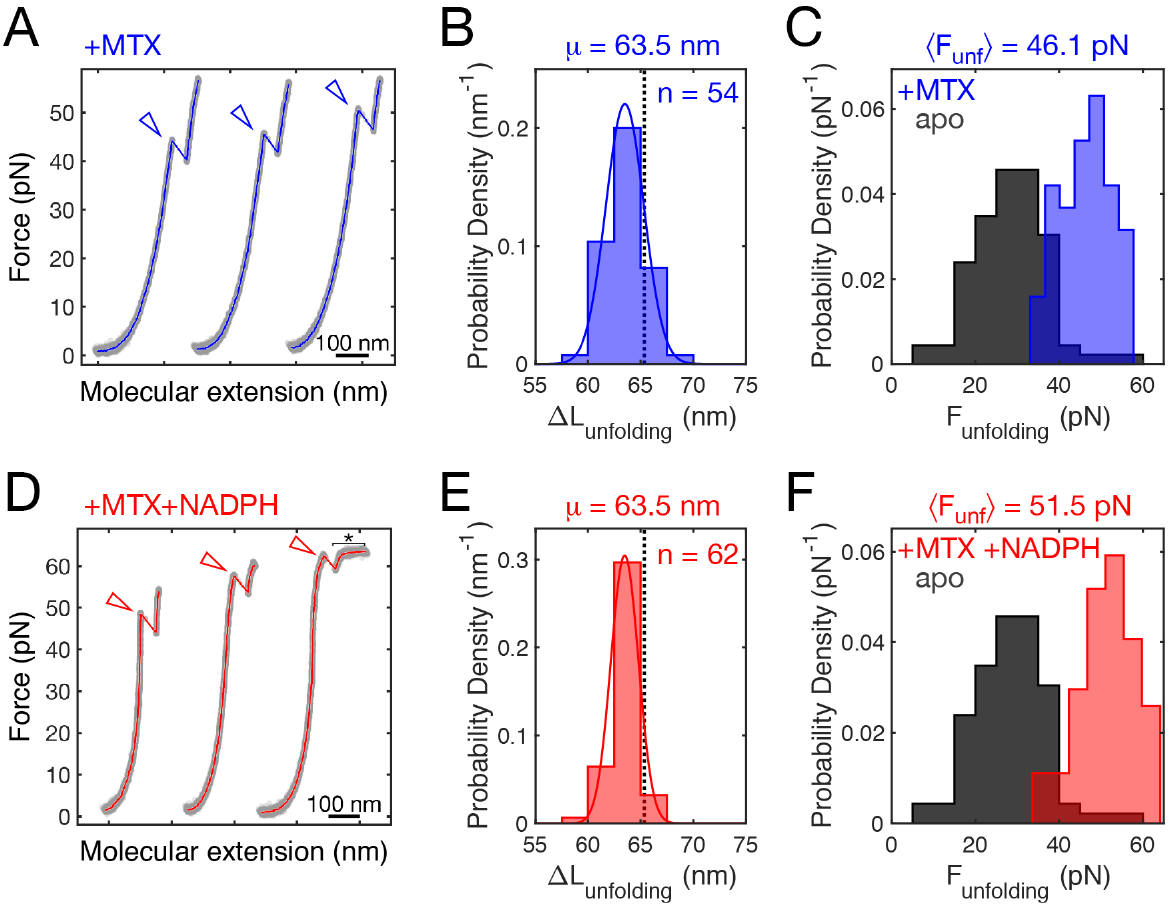
Ligands stabilize mDHFR mechanically. **A, D.** Example force extension curves of initial mDHFR unfolding events (arrowheads) in the presence of MTX (A, blue) and MTX + NADPH (D, red). Unfolding occurs mostly above 40 pN. When both ligands are present, some of the unfolding events occur in the force range of DNA overstretching (D, asterisk). **B, E.** Contour length changes for initial unfolding events from 54 molecules and 62 molecules of either MTX only (B) or MTX + NADPH (E). The means (μ) of normal distribution fits (solid lines) to the data are very similar to each other and close to the calculated value of 65.4 nm (dotted lines). **C, F.** Unfolding force histograms for the two ligand conditions. Data obtained for apo-mDHFR (dark grey; same as in Figure 2F) plotted for reference. The ligands additively stabilize mDHFR. The narrower unfolding force distributions are consistent with unfolding from single states in the two ligand conditions. The average unfolding forces (〈F_unf_〉) are indicated on top of the panels.

Adding both MTX and NADPH further stabilizes mDHFR. Some unfolding events occur near the characteristic overstretching plateau of the DNA handles (Figure 3D). While the distribution of contour length changes is indistinguishable from the MTX condition (Figure 3E), the unfolding forces are shifted toward higher values (Figure 3F) that range from 33 to 64 pN. Taken together, our single-molecule experiments reveal that the ligands MTX and NADPH additively stabilize mDHFR against mechanical denaturation and appear to lock mDHFR in mechanically very stable conformations.

### SecA actively promotes unfolding

Our single-molecule measurement demonstrate that ligand binding stabilizes mDHFR (Figures 2 and 3), which is reflected in slower unfolding of the protein during translocation (Figure 1). Spontaneous unfolding of mDHFR (as posited by the Brownian ratchet model of translocon activity) or active unfolding (which would indicate that SecA functions as a power stroke motor) could both explain this observation. To distinguish between these two scenarios, we quantitatively compared unfolding rates from biochemical translocation experiments to the mechanical properties of mDHFR from optical tweezers measurements.

To define the unfolding kinetics of mDHFR under force, we utilized the method of Dudko and co-workers^48^ to convert the unfolding force distributions into native state lifetimes. Their dependence on force was modeled with an approximation^49^ of Kramers’ theory^50^, which yields an analytical description for the unfolding rates of mDHFR under mechanical load using the parameters k_0_ (the intrinsic unfolding rate), Δx^‡^ (the transition state distance), ΔG^‡^ (the barrier height). This analytical description makes it possible to obtain unfolding rates in force ranges that are not directly accessible in pulling experiments.

We developed a maximum-likelihood method for analyzing unfolding force distributions (see Materials and Methods, and Supplementary Information). To extend the unfolding force range covered in our experiments, we collected data at a lower trap velocity of 20 nm/s. As expected, reduced loading rates result in unfolding at lower forces (Figure 4A and B, histograms). Globally fitting the Kramers-like model to both the slow and fast pulling rate datasets yielded one set of parameters for each ligand condition (Table 2). The force distributions reconstructed from the fitting parameters match the data reasonably well (Figure 4A and B, lines), indicating that our approach yields a good description of the mechanical unfolding kinetics of mDHFR over a relatively wide force range.

**Figure 4.**
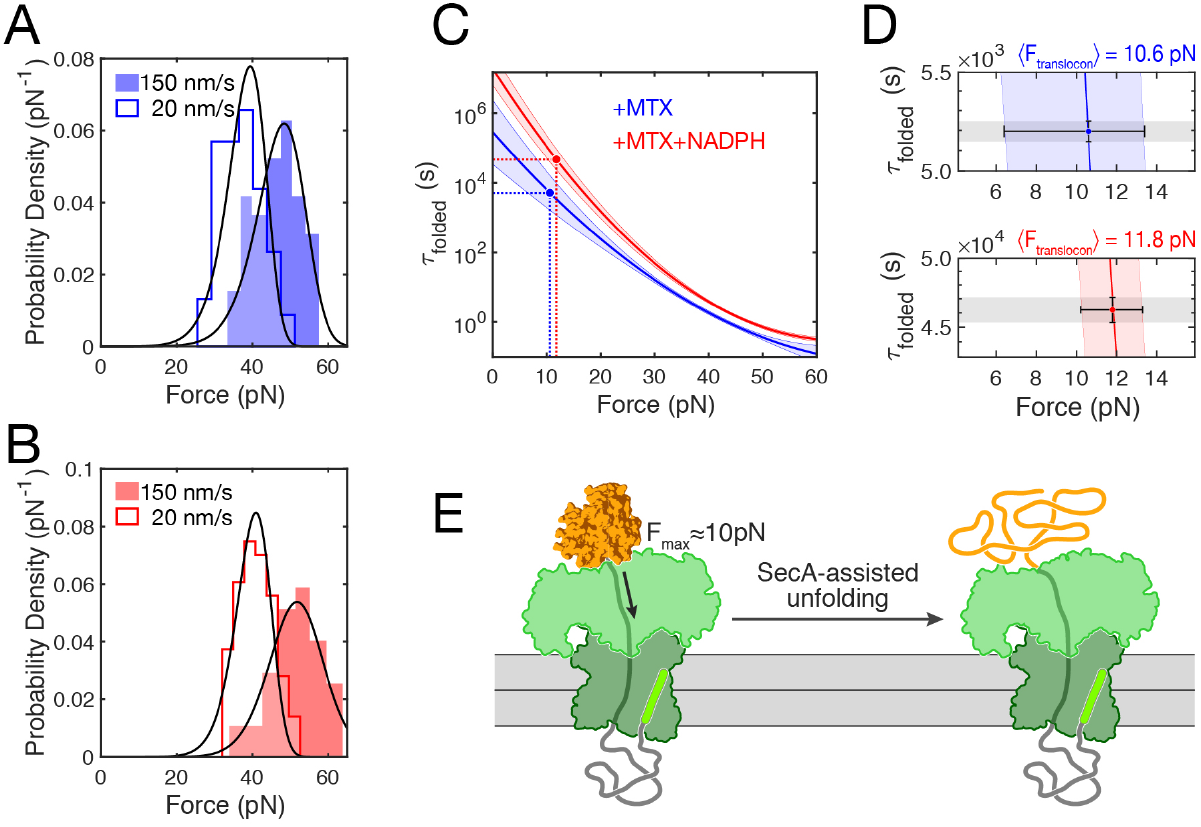
The translocon generates mechanical force to unfold mDHFR. **A, B.** Histograms of unfolding force distributions for single- (A) and double- (B) liganded mDHFR at two pulling speeds. The distributions at a trap velocity of 150 nm/s (filled) are the same as in Figure 3. At a lower trap velocity of 20 nm/s, unfolding occurs on average at lower forces. Solid lines represent probability densities reconstructed using a Kramers-like model with parameters obtained from maximum likelihood analysis of the transformed experimental data. **C.** Dependence of the folded state lifetime of mDHFR (τ_folded_) on force in the presence of MTX (blue) and MTX + NADPH (red). Solid lines are the means obtained from fitting the unfolding force distributions, the shaded regions represent the random error (standard deviation). Solid circles indicate the folded state lifetimes from translocation experiments. In both ligand conditions, similar values for the corresponding forces are obtained (dashed lines). **D.** Close-up of the folded-state lifetimes. Colored lines and shaded areas as in C. The grey shaded area and vertical error bars represent the uncertainty in folded state lifetimes from translocation measurements. The horizontal error bars represent the uncertainty in the translocon force (from standard deviations), obtained by relating the translocation and single-molecule force spectroscopy results. **E.** Schematic illustrating the direction and magnitude of force applied by SecA during translocation of proOmpA-mDHFR.

**Table 2.** Rates from analyzing single-molecule unfolding measurements using Kramers’ theory. The table shows the mean ± standard deviation for the distance to the transition state, barrier height, and intrinsic folded state lifetime (Δx^‡^, ΔG^‡^, τ_o_).

| condition | $\Delta x^\ddagger$ (nm) | $\Delta G^\ddagger$ ($k_B \cdot T$) | $\tau_0$ (s) |
| --- | --- | --- | --- |
| MTX | $1.67 \pm 0.44$ | $15.3 \pm 1.1$ | $2.85 \cdot 10^5 \pm 1.34 \cdot 10^5$ |
| M+N | $2.29 \pm 0.16$ | $18.0 \pm 1.2$ | $1.90 \cdot 10^7 \pm 1.75 \cdot 10^7$ |

The folded state lifetimes at any given force are higher in the double-ligand condition than in the single-ligand case (Figure 4C), reflecting the higher average unfolding forces. Having defined the response of mDHFR to force quantitatively, we can relate it to the unfolding rates observed during translocation to estimate the magnitude of mechanical force that might be applied by the translocon. Matching the respective rates to the force-dependent lifetimes, we obtain similar force values of Ftranslocon(MTX) = 10.6 pN and Ftranslocon(M+N) = 11.8 pN (Figure 4C and D). These results strongly suggest that SecA acts as an active unfoldase, applying the equivalent of approximately 10 pN of constant force to the translocating substrate protein (Figure 4E). Taken together, our data support a model of SecA acting as a power-stroke motor that unfolds proteins during translocation.

## Discussion

A wealth of genetic, biochemical and structural studies of the Sec translocation machinery has yielded a comprehensive understanding of its constituent parts and their activities. Mechanical aspects have been proposed to be important for Sec translocon function^7^’^9,14^, but their experimental investigation is challenging. Here, we have mechanically calibrated a widely employed probe for protein transport studies, mDHFR, using optical tweezers (Figures 2 and 3). Analyzing the Sec-dependent translocation kinetics of this probe by developing a comprehensive model (Figure 1) has allowed us to estimate for the first time, to our knowledge, the magnitude of mechanical force that a folded substrate protein is subjected to during the process (Figure 4).

Our results indicate that the reconstituted minimal Sec machinery (SecYEG and SecA) can generate mechanical force to the translocating polypeptide that is equivalent to a constant tension of approximately 10 pN. A similar value was deduced from analyzing the effective force needed to overcome regulatory translation elongation arrest by the SecM protein^51^. The dedicated protein unfoldase ClpX was found to generate up to 20 pN of mechanical force during regulated protein degradation^17^ and thus operates in a similar force regime. ClpX, acting together with the protease ClpP, enables efficient degradation of a large variety of stably folded intracellular proteins. The force determined here for SecA should therefore be sufficient to ensure reliable translocation through roadblocks in translocation substrates.

mDHFR is a widely utilized force probe in protein transport experiments^7,23–27^. As such, a thorough mechanical characterization of its mechanical response is essential for quantitatively interpreting such measurements. Our single-molecule experiments suggest that apo-mDHFR populates several native states of distinct mechanical stabilities (Figure 2). The bacterial ortholog, whose folding has been characterized in detail^47^, was found to exist in several interconverting native states. A similar conformational equilibrium, in which the protein switches between states that have distinct mechanical stabilities, is consistent with the force distribution observed upon initial unfolding of mDHFR in our experiments (Figure 2F). We also find that the protein does not efficiently refold within 15 seconds at a low force of 2 pN (Figure 2D). Because the N-terminus of the unfolded protein is expected to be quickly sequestered by the translocon, refolding of mDHFR is not expected to occur during translocation. Initial unfolding therefore likely represents the rate-limiting step.

The ligands MTX and NADPH additively stabilize mDHFR mechanically (Figure 3). The ligand-bound protein does not exhibit the apparent heterogeneity observed for the apo-state. MTX and NADPH might lock mDHFR in a structure of high mechanical resistance, preventing conformational excursions to more labile states. Mechanical stabilization of native mDHFR was not observed in atomic force microscopy experiments^45^. This discrepancy is likely due to difference in experimental conditions. The high loading rates in atomic force microscopy experiments might result in different barriers being probed. While the details of force generation by the translocon are unknown, the lower loading rates in optical tweezers experiments are likely closer to those during translocation.

Structural studies^11,52,53^ suggest that tension on the translocating polypeptide arises when the SecA two-helix finger pulls the folded domain against the clamp. The axis of force application depends on how the domain abuts the clamp, which is not known. In our optical tweezers experiments, we apply force to the termini of mDHFR, which are located on the same side of the central beta-sheet (Figure 1A). As a result, unfolding occurs in an “unzipping” geometry, in which force is applied perpendicularly to the strands (Figure 2A). This configuration is mechanically more labile than a “shearing” geometry^54,55^, in which force is applied to opposite sides of the betasheet. In pulling experiments with titin I27 domain, such a shearing geometry gives rise to very high unfolding forces^54^ that are unlikely to be generated by the Sec system. The observation that the translocon unfolds the I27 domain^14^ therefore suggests an unzipping geometry. For mDHFR, the unfolding geometries in translocation and single-molecule pulling experiments are likely more similar to each other. Nevertheless, the pure unzipping geometry in our optical tweezers measurements of mDHFR unfolding might underestimate the magnitude of the mechanical load generated by SecA.

Biochemical experiments^56^ show that ADP release is the rate-limiting step during the SecA ATPase cycle. As a consequence, the translocon resides in the ADP-bound state for much of the ATP hydrolysis cycle. smFRET experiments suggest that the substrate protein is free to slide through the translocon in the ADP state, with the two-helix finger disengaged and the clamp open^9^. As such, force is not continuously applied. Rather, the polypeptide experiences force during less than half of the ATPase cycle and may slide freely during the remaining time^9^. It therefore seems possible that the actual force applied by SecA to achieve substrate unfolding at the observed rates is higher than our analyses suggest.

Our single-molecule experiments reveal that ligand-bound mDHFR is highly stable, exhibiting very low zero-force unfolding rates (on the order of ~10^−6^ s^−1^, Figure 4C). Rectifying spontaneous structural transitions, as postulated in a Brownian ratchet mechanism of SecA action, would therefore result in overall translocation rates for ligand-bound mDHFR much slower than those observed in our experiments. A “steric force”, similar to the one generated by a nascent protein on the ribosome^51^, could in principle accelerate unfolding of a translocating polypeptide held in close proximity to the translocon by a passive ratchet. However, a passively sliding substrate is statistically unlikely to assume a position close to enough to the translocon for a significant steric force to arise. An active force-generating power-stroke model of SecA function therefore better explains our observations.

Taken together, our experiments indicate that the SecA motor generates mechanical forces of 10 pN or more, comparable to the dedicated protein unfoldase ClpX, and that it likely acts as a power-stroke motor. The ability to actively unfold tertiary structure in substrate proteins might help to safeguard the translocation system against jamming by substrate proteins that escape holdase chaperones and fold before passing through the translocon channel. The exact mechanisms by which SecA converts chemical energy into mechanical work remain to be determined. Our analyses are an important step toward elucidating the mechanochemistry of protein translocation by the Sec translocon.

## Materials and Methods

### Cloning, expression and purification of pOA-mDHFR

The coding sequence for the first 178 amino acids of pOA were amplified by PCR from a parent pOA plasmid using primers for Gibson assembly that added a GSGS linker at the C-terminus. The entire mDHFR ORF was PCR amplified from a mDHFR plasmid with a C-terminal AviTag fused to the mDHFR fragment using Gibson Assembly (NEB). The backbone contained a T7 promoter. Cysteines were added at the desired positions using either site directed mutagenesis or by insertion of synthetic DNA fragments (gBlocks, IDT DNA). The p86 peptide sequence VSGWRLFKKIS^32^ was added to the end of the of the construct via PCR. The resulting plasmid (termed pOA-mDHFR) was transformed into *E. coli* strain MM52, which contains temperature sensitive SecA allele. A starter culture was grown at the permissive temperature (30°C) until it reached OD_600_ of 0.5. The culture was then shifted to the restrictive temperature (37°C), and expression was induced with 1 mM IPTG for four hours. The cell pellet was harvested and washed with cold 1X PBS. Cells were lysed using an EmulsiFlex-C5 (Avestin) in 1X PBS pH 7.5. Inclusion bodies were washed with 1X PBS three times. Inclusion bodies were pelleted and frozen in liquid N2. The substrate protein was solubilized from inclusion bodies in 6M urea before use.

### Expression and purification of SecYEG

Plasmid pSOS334, encoding cysteine free His-SecY, SecE and SecG, was a generous gift from Dr. Shu-ou Shan. Transformed BL21-Gold(DE3) cells were grown to an OD_600_ of 0.4 to 0.6 at 37°C before expression was induced by adding IPTG to a final concentration of 0.5 mM IPTG. Cells were harvested 2 hours after induction at 37°C. Cells were harvested and lysed by sonication. After centrifugation at 30,000 g to remove cell debris, membranes were isolated by ultracentrifugation and solubilized in dodecyl-β-maltoside (DDM). SecYEG was isolated by ion exchange chromatography on a sulfo-propyl resin, followed by affinity purification with NiNTA resin. The purified protein was flash-frozen and stored at −80 °C in 50 mM HEPES-KOH at pH 7.5, 150 mM KOAc, 20% (w/v) glycerol, and 0.2% DDM.

### Expression and purification of SecA

A plasmid encoding cysteine-free SecA with an N-terminal His tag was transformed into BL21(DE3) cells. Protein expression was induced with final concentration of 0.5 mM IPTG at 37°C in cultures with an OD_600_ of 0.4 to 0.6. Cells were harvested 2 hours after induction at 37°C. Cells were lysed by sonication in 50 mM HEPES-KOH pH 7.6, 150 mM KOAc, 10% glycerol with 1 tablet protease inhibitor cocktail (Roche). After cell lysis and centrifugation at 30,000*g*, 4°C to remove cell debris, the protein was purified from the supernatant by affinity chromatography on a NiNTA resin in 50 mM HEPES-KOH pH 7.6, 150 mM KOAC, 10 % glycerol, and eluted with 300 mM imidazole. The eluted product was dialyzed overnight against the same buffer without imidazole. Protein aliquots were flash-frozen and stored at −80°C.

### Cloning, Expression and Purification of NanoLuc 11S and GST-dark

The codon optimized 11S sequence was obtained as a synthetic DNA fragment (IDT DNA) and cloned into a His-SUMO backbone (Addgene Plasmid #37507) for expression. The coding sequence for the p86 **“**dark” peptide (VSGWALFKKIS)^31^ was inserted into a plasmid encoding glutathione S transferase (GST) to generate a C-terminal fusion in the same backbone and termed “GST-dark.” For expression, plasmids were transformed into BL21(DE3) cells. Expression was induced with final concentration of 0.2% arabinose at 37°C when OD_600_ reached 0.4**~**0.6. Cells were harvested 4 hours after induction at 37°C and lysed by sonication in 1x PBS pH 7.5 with 1 tablet protease inhibitor cocktail (Roche). After cell lysis and centrifugation at 30,000*g*, 4°C to remove cell debris, the protein was affinity purified from the supernatant using a 5 mL HisTrap column (GE Healthcare) in 1XPBS pH 7.5 and eluted with 300 mM imidazole. After cleavage of the His-SUMO tag, pure protein was obtained by reverse Ni-NTA affinity chromatography in 1x PBS pH 7.5. The protein was concentrated, aliquoted, and flash frozen in liquid N2.

### Encapsulation of 11S in SecYEG/SecA proteoliposomes

Unilamellar liposomes were prepared by extrusion of *E. coli* polar lipids (Avanti) suspended in 10 mM Hepes at pH 7.5, 100 mM NaCl through membranes with a 200-nm pore diameter. In order to swell the liposomes, 4.7 mM DDM was added to 5 mM lipids. After incubation at room temperature for 3 hours, proteins (5 μM SecYEG, 5 μM SecA, and 50 μM 11S) were added to liposomes. The reaction was incubated for 1 hour at 4°C, followed by 4 incubations with BioBeads SM-2 (BioRad) to remove the detergent. The proteoliposomes were isolated by centrifugation at 250,000*g*, 30 minutes at 4 °C, in a TL100 rotor (Beckman). The pellet was resuspended in 10 mM Hepes at pH 7.5, 100 mM NaCl and run over a Sephacryl 200 column in order to remove unincorporated 11S from the proteoliposomes. Proteoliposomes were recovered by ultracentrifugation before resuspension. They were flash frozen in liquid N_2_ for storage at −80°C.

### Intramolecular Disulfide Bond formation in pOA-mDHFR

Intramolecular disulfide bond formation in pOA-mDHFR was achieved by incubation with Cu^2+^/phenanthroline^57^. 1 mM Cu^2+^/phenanthroline was added to 10 μM protein and incubated at 4°C for 18 hours. Disulfide bond formation was assessed by SDS-PAGE. Oxidized protein was flash frozen in liquid N2 and stored at −80°C until needed for translocation experiments.

### Protease-protection translocation assay

Translocation substrates were synthesized *in vitro* using the PURExpress system (NEB) in the presence of [^35^S]-methionine for 2 hours at 37°C. The translation reaction was transferred to ice and precipitated with three volumes of saturated ammonium sulfate in 20 mM HEPES. Precipitated protein was pelleted at 14000 rpm for 15 minutes. The pellet was resuspended in 6 M urea, 20 mM Tris-HCl, pH 6.8. Disulfide bond formation was achieved by incubation with 400 μM sodium tetrathionate for 30 minutes. Translocation reactions were performed in the presence of 0.2 μM SecYEG in proteoliposomes, 1.2μM SecA, and in vitro synthesized substrate diluted 1:50, and in 50 mM phosphate buffer, pH 7.5, 20 μg/mL BSA. After addition of 5 mM ATP, translocation up to the disulfide bond was allowed to proceed for 15 minutes. Subsequently, the disulfide bond in the substrate was reduced with 50 mM DTT, marking time zero. After translocation at 37°C, sample were taken at defined time points and the reaction was quenched by addition of 18 mM EDTA and 1.5 M urea. Samples were then treated with 2 mg/mL proteinase K on ice for 30 minutes, precipitated with 10% trichloroacetic acid, and analyzed by SDS-PAGE and autoradiography.

### Real-time translocation assay

11S encapsulated proteoliposomes were incubated with buffer (10 mM HEPES, pH 8, 100 mM KOAc, 5 mM Mg(OAc)2) and 1 μM GST-dark for 10 minutes at room temperature. pOA-mDHFR containing a disulfide bond was added along with NanoGlo Live Cell Assay Buffer with NanoGlo substrate. After the addition of 5 mM ATP the reaction was taken to a plate reader (Promega GloMax Navigator) and the luminescence collected for approximately 30 minutes. During this time, additional GST-dark (0.5 μM) was added. After the luminescence signal plateaued, 1 mM DTT was added to reduce the disulfide bond and restart translocation, followed by luminescence data collection for 30 minutes. Luminescence readings were collected continuously at a sampling rate of 0.2 Hz.

### Cloning, expression, and purification of mDHFR for optical tweezers experiments

To generate an expression construct for mDHFR, we amplified the open reading frame from a mouse cDNA library using polymerase chain reaction and inserted it into a pBAD His6 Sumo TEV LIC cloning vector (Addgene Plasmid #37507) that had been engineered to encode an N-terminal Avi tag and C-terminal ybbR tag^58^. The mDHFR plasmid was transformed into BL21-Gold(DE3) (Agilent Technologies) host cells, and protein expression was induced with final concentration of 0.2% (w/v) L-arabinose (AMRESCO) at 37°C when OD_600_ reached 0.4**~**0.6. Cells were harvested 4 h after induction at 37°C. Cells were lysed using an EmulsiFlex-C5 homogenizer (Avestin) in 1X PBS pH 7.5 with two tablets of CompleteMini EDTA-free protease inhibitor (Roche). After cell lysis and centrifugation at 30,000*g*, 4°C to remove cell debris, the protein was affinity purified from the supernatant using a 5 mL HisTrap column (GE Healthcare). The protein was dialyzed against 1X PBS overnight in the presence of 1:1000 (w/w) Ulp1 to remove the His6-SUMO tag. The cleaved protein was applied to the HisTrap column again to remove the His6-SUMO moiety and the His6-tagged Ulp1 enzyme. The purified protein in the flow through was then incubated with 1 μM BirA biotin ligase in 1X biotinylation buffer (25 μM D-biotin, 5 mM ATP, and 5 mM Mg-acetate) at 25°C for 1 h to ensure complete biotinylation of the Avi-tagged protein. Insoluble protein was removed via centrifugation at 14,000 rpm, 4°C. The product was concentrated and loaded onto a 5 mL HisTrap column, and the flow through collected. After concentration, protein aliquots were flash-frozen and stored at −80°C.

### Derivitization of mDHFR

In order to immobilize mDHFR on polystyrene beads for optical tweezers experiments, we modified the biotinylated, ybbR-tagged protein with a CoA-modified double-stranded DNA (dsOligo-CoA) that also contained a “sticky end” for ligation in an Sfp-mediated reaction^58^. After the reaction, the sample was centrifuged briefly (10 min, 16,000*g*, 4°C) and loaded onto a Superdex 200 column (GE Healthcare) to remove Sfp and unreacted dsOligo-CoA. Successful derivatization was confirmed using SDS-PAGE, and the modified protein was flash frozen in small aliquots and stored at −80°C.

### Optical tweezers experiments

We carried out optical tweezers experiments using a single trap optical tweezers instrument with two counter-propagating 845 nm diode lasers^59^. Single mDHFR molecules were tethered as described by Liu *et al*.^58^. All optical tweezers experiments were performed in 1X PBS, 1 mM EDTA and ligand depending on the experimental condition. One bead was held in the optical trap while the other was held on the micropipette. Force ramp data were collected with pulling velocity of 20 nm/s or 150 nm/s and a trap stiffness of ~0.1 pN/nm. The force was increased until an unfolding event was observed, after which the force was decreased to 2 pN and the protein was allowed to refold for 15 seconds. Unfolding forces and extension changes were determined as described in detail in Liu *et al*.^58^. Contour length changes were calculated from extension changes using a worm-like chain model with a persistence length of 0.65 nm. Expected contour length changes were calculated using a contour length increment of 0.36 nm per amino acid for the unfolded polypeptide and a native-state end-to-end distance of 1.6 nm, determined from the mDHFR crystal structure coordinates pdb 1u70.

### Determination of kinetic rates from translocation data

The time evolution of the luminescence signal in translocation experiments reflects the kinetics of sequential processes that result in substrate protein import, as well as interfering side-processes. The productive processes considered in our mathematical model are translocation, stalling at the disulfide loop, and unfolding. A competing process results in translocon incapacitation, i.e. the irreversible loss of translocation activity. Our model quantitatively describes translocation of pOA-mDHFR as a series of events: translocon engagement, stalling at the disulfide loop, translocation to the folded mDHFR roadblock, mDHFR unfolding and completion of translocation. We also take into account complicating processes that contribute to the total signal, namely the presence of substrate without a disulfide loop, incapacitation, and luciferase substrate depletion. Supplementary Figure 2 provides a graphical overview of the model. A full description of the mathematical model is provided in Supplementary Information.

### Determination of unfolding rates from force spectroscopy data

The protein unfolding rate in force spectroscopy measurements depends on the applied force. This force dependence is reflected in the distribution of rip forces from force ramp experiments. Dudko *et al*.^48^ developed a method to transform unfolding force distributions into force-dependent lifetimes. The force-dependent lifetimes can then be fit with an appropriate model to obtain an analytical expression for the unfolding rate as a function of force. This approach typically requires binning of the unfolding forces, resulting in loss of force resolution due to finite bin size. To avoid this information loss, we developed an analysis procedure that converts the force dependence of the unfolding rate into a probability distribution, which we then fit to our data using the method of maximum likelihood. This approach allowed us to use the experimentally observed loading rate for each molecule individually and calculate a global fit for data sets that cover a wide range of loading rates. A full description of the analysis method is provided in Supplementary Information.

## Acknowledgements

R.G. acknowledges support from the CMDB graduate program (NIH 5T32GM007231). C.M.K. acknowledges support from the NIH (5R01GM121567 and 5R21AI133514). We are grateful to Dr. Shu-ou Shan (Caltech) for the gift of the SecYEG expression plasmid, Dr. Ian Collinson (University of Bristol) for advice regarding the translocation system, and Dr. Donald Oliver (Wesleyan University) for the gift of *E. coli* strain MM52. We thank members of the Kaiser lab for helpful discussions.

