## Supplementary Information for "The SecA motor generates mechanical force during protein translocation"

***The SecA motor generates mechanical force to unfold translocating proteins***

Supplementary information contains

- Supplementary Figures S1 to S5
- Supplementary Table ST1
- Supplementary Methods

#### Supplementary Figures

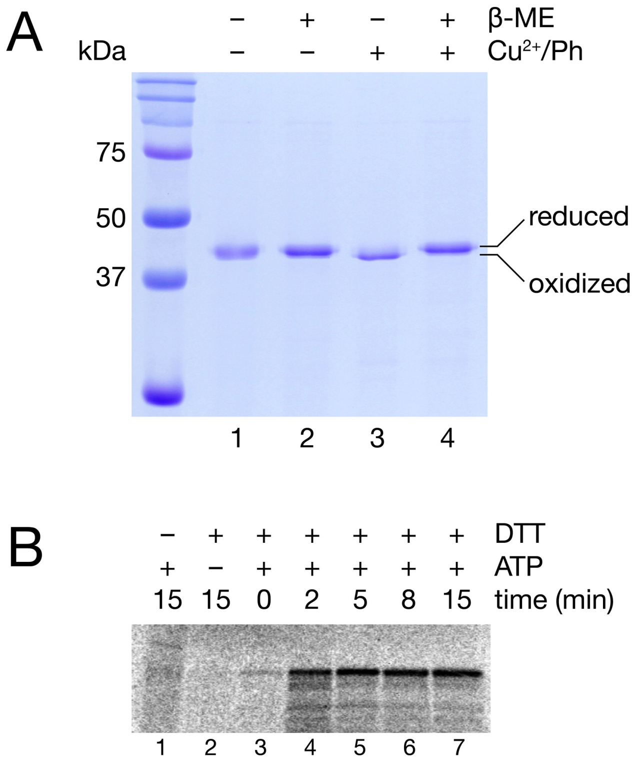


**Supplementary Figure S1. Disulfide loop formation reversibly stalls translocation. A.** Loop formation of pOA-mDHFR. pOA-mDHFR was incubated with copper/phenanthroline ($Cu^{2+}$/Ph) to catalyze the formation of an intramolecular disulfide bond. Oxidized pOA-mDHFR migrates with higher electrophoretic mobility than the reduced form, which is expected for a polypeptide containing a disulfide loop (lane 3 vs. lane 4). **B.** Translocation of the pOA-mDHFR substrate protein assessed by protease protection. Radiolabeled oxidized protein was added to SecYEG/SecA proteoliposomes and incubated at 37°C for the time indicated. The sample was then subjected to proteinase K digestion, and products were analyzed by SDS-PAGE and autoradiography. In the absence of DTT or ATP, little protected substrate protein is observed (lanes 1 and 2). When both ATP and DTT are added, an increasing amount of protected protein is observed (lanes 3 to 7), indicating translocation into the interior of the proteoliposomes.

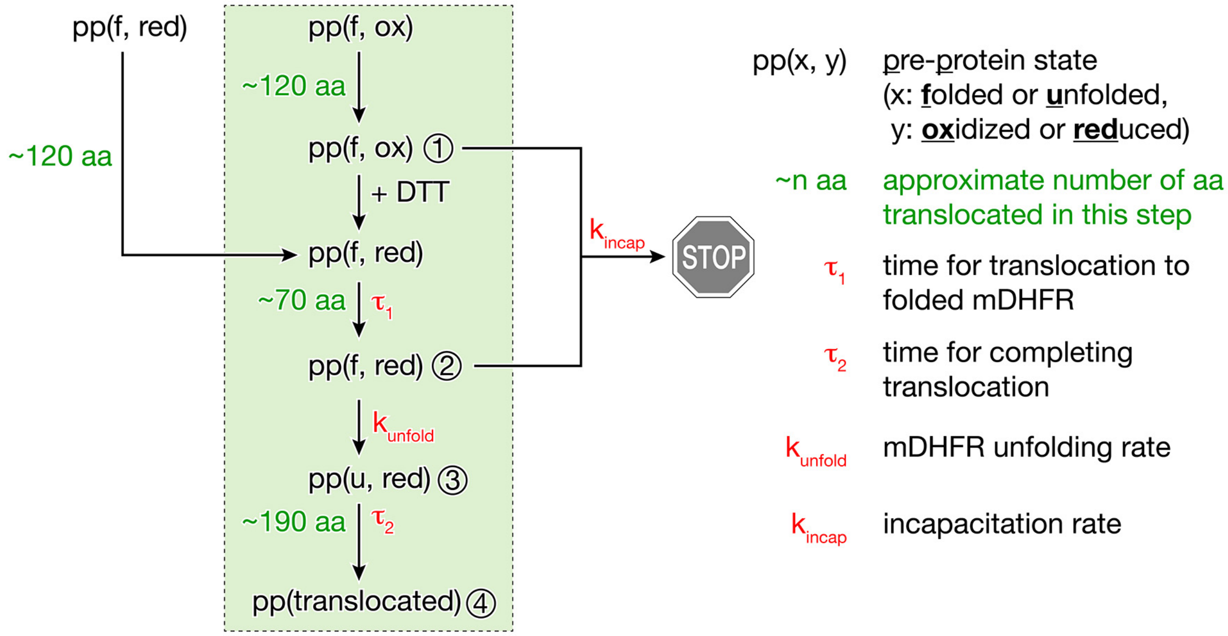


**Supplementary Figure S2. Schematic of our kinetic model of translocation.** The luminescence signal readings from translocation experiments reflect the kinetics of multiple processes. The processes relevant to our analyses are shown in the green box. At the beginning of the measurement, the preprotein (pp) is primarily in an oxidized and folded state (pp(f,ox)). In the presence of ATP, translocation of the first ~120 amino acids proceeds until stalling at the disulfide loop (①). DTT is added after a waiting time that allows accumulation of stalled substrate protein, yielding reduced preprotein (pp(f,red)). Translocation of the next ~70 amino acids proceeds with the time constant $\tau_1$ until the translocon encounters folded mDHFR (②). After unfolding of mDHFR with the rate $k_{unfold}$, the unfolded preprotein (pp(u, red)) (③) completes translocation with the time constant $\tau_2$, resulting in translocated preprotein (pp(translocated)) inside the proteoliposomes (④). The model accounts for the presence of reduced preprotein that undergoes translocation independently of DTT addition (left). An irreversible side process that we term incapacitation (right) occurs with rate $k_{incap}$, resulting in reduced amounts of translocated protein. Additional parameters in the model include the depletion of the luciferase small molecule substrate and instrument sensitivity, which are not shown in this diagram. The fit parameters that yield the translocation and unfolding rates are shown in red here. See Supplementary Information for a detailed description of the model.

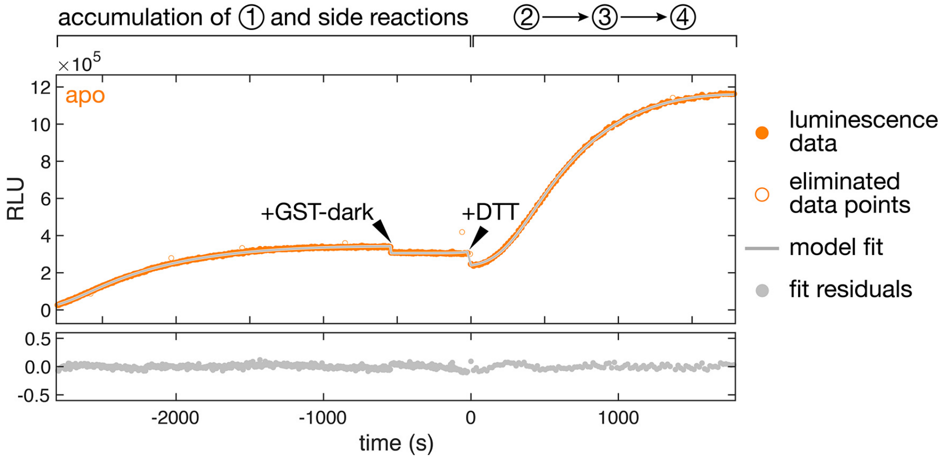


**Supplementary Figure S3. Real-time translocation measurement.** Complete luminescence recording (upper graph) for a translocation experiment with pOA-mDHFR in the absence of ligands starting after mixing SecYEG/SecA proteoliposomes containing encapsulated 11S NanoLuc, ATP, $GST_{dark}$ and oxidized pOA-mDHFR substrate protein. Closed orange circles represent the data used to calculate a fit (grey line) based on our model. Open circles represent data points that were eliminated from analysis because they had an unusually high variance (9 out of 566 samples in this recording; see Supplementary Information, section 1, for details). The initial increase results from side reactions (presumably import of non-oxidized substrate protein molecules) that occur while stalled translocation substrate accumulates. Arrowheads indicate the addition of extra GST-dark and DTT to the reaction. Additional GST-dark quenches any 11S protein that is accessible on the outside of the proteoliposomes. DTT reduces the substrate protein, restarting translocation. The reaction stages are indicated on top, using the same numbers as in Figure 1C. The graph on the bottom shows the residuals from the fit.

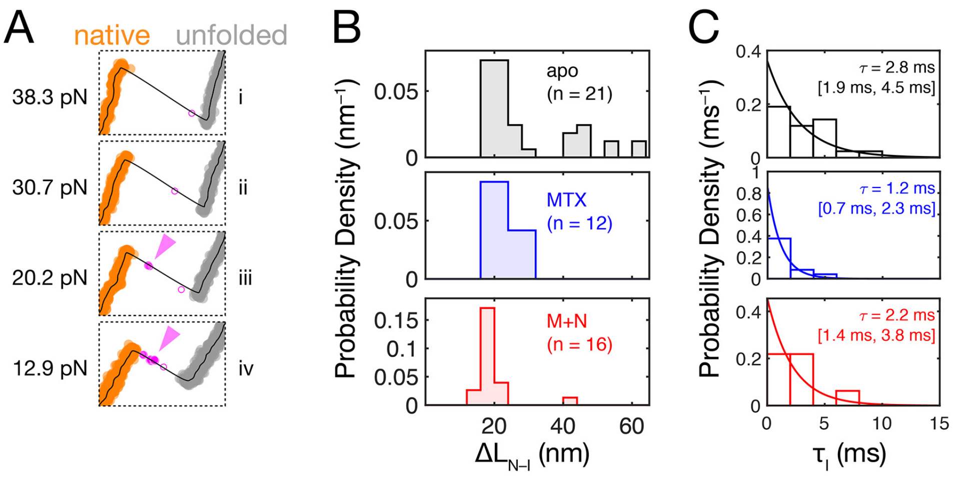


**Supplementary Figure S4. mDHFR populates a transient unfolding intermediate. A.** Blow-ups of the region around the unfolding transition for four apo-mDHFR molecules. Raw data is shown as circles, colored by state (orange: folded, grey: unfolded), with samples that fall between the two states in magenta. One or two samples between folded and unfolded can result from time-averaging (white with magenta outline). The majority of traces do not exhibit a detectable intermediate (examples i and ii). Additional samples in the transition region (solid magenta) represent a transient unfolding intermediate, which is apparent in some traces (examples iii and iv, arrowheads). **B.** Histograms of the contour length changes for native-to-intermediate transitions for the three ligand conditions (apo, MTX, M+N). Most transitions result in a contour length change of ~20 nm. The number of traces with a detected unfolding intermediate is indicated in parentheses. **C.** Histograms of intermediate lifetimes (bars) with fit to an exponential probability density function (line). Almost all detected intermediates unfold within 10 ms. The fits yield mean lifetimes of approximately 2 ms. However, our time resolution of 1 ms results in many missed events, and the true intermediate lifetime may be smaller than this value. The lifetimes do not appear to increase in the presence of ligands.

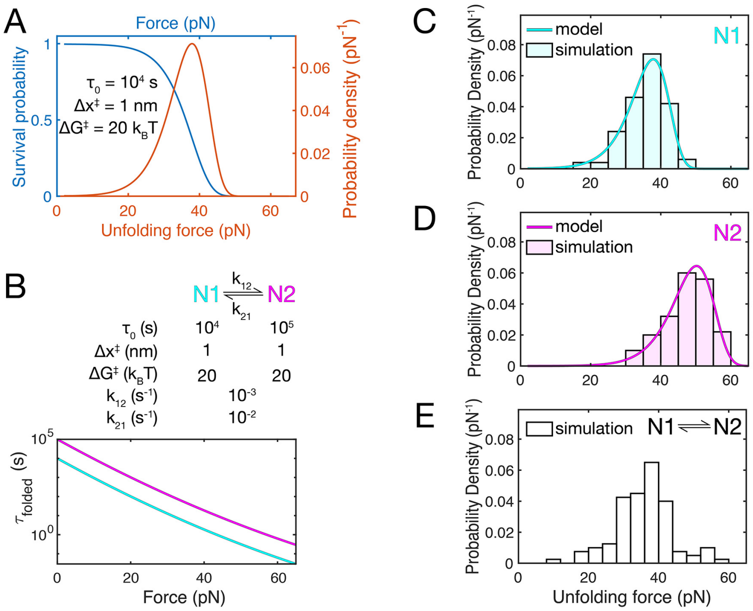


**Supplementary Figure S5. Unfolding force distributions of a state-switching protein in force ramp experiments. A.** Calculated survival probability of the folded state (blue) and unfolding force probability density (orange) in a force ramp experiment using the parameters indicated in the figure. The probability of the folded state staying intact decreases as the force is ramped up (blue line). As a result, the probability density of the observed unfolding forces (orange line) shows a skewed distribution that drops sharply at high forces. **B.** Parameters for a state switching model and calculated force-dependent lifetimes. In the hypothetical model, two states (N1 and N2) that differ in their intrinsic folded state lifetimes interconvert with constant rates ($k_{12}$ and $k_{21}$). **C, D.** Calculated (lines) and simulated (bars) lifetimes distributions for states N1 and N2. The simulated data (100 events) shows the expected distributions. **E.** Simulated data for a state-switching model. The simulated unfolding forces (100 events) show a broad distribution with a tail at high unfolding forces, similar to the experimentally observed distribution for mDHFR (Figure 2F).

#### Supplementary Table

| | apo | | MTX | | MTX+NADPH | |
|---|---|---|---|---|---|---|
| **Parameter** | **mean** | **sd** | **mean** | **sd** | **mean** | **sd** |
| $\alpha_1$ | $4.19 \cdot 10^{-4}\ s^{-1}$ | $2.56 \cdot 10^{-6}\ s^{-1}$ | $1.92 \cdot 10^{-4}\ s^{-1}$ | $1.89 \cdot 10^{-6}\ s^{-1}$ | $2.16 \cdot 10^{-5}\ s^{-1}$ | $4.16 \cdot 10^{-7}\ s^{-1}$ |
| $\alpha_2$ | $1.10 \cdot 10^{-3}\ s^{-1}$ | $1.78 \cdot 10^{-6}\ s^{-1}$ | $1.21 \cdot 10^{-3}\ s^{-1}$ | $3.12 \cdot 10^{-6}\ s^{-1}$ | $9.12 \cdot 10^{-4}\ s^{-1}$ | $6.86 \cdot 10^{-6}\ s^{-1}$ |
| $\alpha_3$ | 0.985 | $6.61 \cdot 10^{-5}$ | 0.982 | $1.42 \cdot 10^{-4}$ | 0.897 | $1.62 \cdot 10^{-3}$ |
| $\alpha_4$ | 245 s | 0.588 s | 251 s | 0.864 s | 261 s | 2.19 s |
| $\alpha_5$ | 148 s | 1.10 s | 156 s | 1.68 s | 157 s | 4.89 s |
| $\alpha_6$ | 16.1 s | 0.752 s | 75.8 s | 0.991 s | 116 s | 2.23 s |
| $\alpha_7$ | 132 s | 1.70 s | 130 s | 2.01 s | 296 s | 2.64 s |
| $\alpha_8$ | $6.28 \cdot 10^{-11}$ | $7.65 \cdot 10^{-13}$ | $6.28 \cdot 10^{-11}$ | $7.65 \cdot 10^{-13}$ | $6.28 \cdot 10^{-11}$ | $7.65 \cdot 10^{-13}$ |
| $\alpha_9$ | $6.37 \cdot 10^{+7}$ | $2.65 \cdot 10^{+6}$ | $6.37 \cdot 10^{+7}$ | $2.65 \cdot 10^{+6}$ | $6.37 \cdot 10^{+7}$ | $2.65 \cdot 10^{+6}$ |
| $\alpha_{10}$ | 0.910 | $9.00 \cdot 10^{-4}$ | 0.921 | $1.20 \cdot 10^{-3}$ | 0.910 | $1.40 \cdot 10^{-3}$ |
| $\alpha_{11}$ | 0.693 | $2.20 \cdot 10^{-3}$ | 0.703 | $2.53 \cdot 10^{-3}$ | 0.921 | $2.46 \cdot 10^{-3}$ |

**Supplementary Table ST1. Full parameter set for fits to real-time translocation measurements.** See Supplementary Information, section "Kinetic Model for Fitting Luminescence Data"  for a detailed description of the model.

### Supplementary Methods

#### 1. Kinetic Model for Fitting Luminescence Data

In our translocation measurements, the luminescence signal is generated when a p86-tagged substrate protein is fully translocated into the interior of SecYEG/SecA proteoliposomes and restores luciferase activity by binding to 11S. We developed a kinetic model that takes into account the following processes:

(1) The oxidized substrate protein engages with the translocon. After DTT-induced disulfide loop opening, unfolding of mDHFR and completion of translocation, light is generated by the encapsulated NanoLuc luciferase.

(2) Substrate protein with reduced cysteines bypasses disulfide-loop stalling and reaches the interior of the proteoliposomes, generating signal.

(3) Irreversible inactivation (“incapacitation”) prevents substrate import.

(4) NanoLuc substrate depletion results in a time-dependent decrease of light intensity.

Our model describes the time evolution of the luminescence intensity, I, as a function

$$I = I(t, \boldsymbol{\alpha}) \qquad \text{Eq. S1}$$

where $t$ is time (the independent variable) and $\boldsymbol{\alpha}$ is a parameter vector with the following components:

- $\alpha_1$: unfolding rate of mDHFR
- $\alpha_2$: incapacitation rate
- $\alpha_3$: molar fraction of preprotein with oxidized disulfide at $t = T_0$
- $\alpha_4$: mean translocation time for oxidized mDHFR
- $\alpha_5$: RMS deviation translocation time for oxidized mDHFR
- $\alpha_6$: mean translocation time for reduced mDHFR
- $\alpha_7$: RMS deviation translocation time for reduced mDHFR
- $\alpha_8$: furimazine substrate depletion rate
- $\alpha_9$: scaling factor for instrument sensitivity
- $\alpha_{10}$: relative change in luminescence reading after dilution (at $T_G$)
- $\alpha_{11}$: relative change in luminescence reading after dilution (at $T_R$)

At $t = T_0$, the experiment is set up by mixing proteoliposomes with substrate and ATP. At $t = T_G$, GST-dark is added to the reaction. At $t = T_R$, DTT is added to the reaction. While $t < T_R$ (i.e. before DTT addition), the oxidized substrate protein engages with the translocon and is translocated until the disulfide loop. We consider the disulfide loop reduction after DTT addition to be essentially instantaneous. Thus, translocation resumes after reduction. The mean total time for substrate import into the interior of the proteoliposomes is the sum of mean time required for translocation ($\alpha_4$) and the mean time required for unfolding ($1/\alpha_1$). The variation in translocation time between molecules is modeled as a Gaussian distribution with standard deviation of $\alpha_5$ around the mean $\alpha_4$. Unfolding is assumed to be a first-order process with rate $\alpha_1$. When the substrate reaches the interior of the proteoliposome, its C-terminal p86 tag restores NanoLuc activity by binding to the encapsulated 11S. Active NanoLuc uses furimazine substrate to generate light. As a consequence, the intra-liposome furimazine concentration decreases, which is captured by parameter $\alpha_8$ as described below.

Not all substrate protein molecules are oxidized at $t = T_0$. As a consequence, full translocation can already occur at $t < T_R$. To account for the signal generated by bypassing disulfide loop stalling, we consider that a fraction $(1 - \alpha_3)$ of the substrate protein is not oxidized. These molecules translocate into the interior of the vesicles (translocation time described by $\alpha_6$ and $\alpha_7$, unfolding described by $\alpha_1$), starting at $t = T_0$, and contribute to the total amount of substrate protein inside proteoliposomes and to the overall luminescence signal.

When translocation is halted (either at the disulfide loop or at the folded mDHFR), irreversible "incapacitation" leads to irreversible loss of translocation activity. The molecular basis for incapacitation is not known. Possible processes include translocon inactivation, and substrate protein aggregation or sequestration. Incapacitation occurs with rate $\alpha_2$ when translocation is stalled, resulting in lower overall signal. Because incapacitation occurs at a constant rate and competes with unfolding, the overall efficiency of translocation is scaled by a factor of $\alpha_1 / (\alpha_1 + \alpha_2)$.

The total time between DTT addition and appearance of the p86 tag inside the liposomes consists of three parts: the time $\tau_1$ required for translocation to folded mDHFR, the time required for mDHFR unfolding, and the time $\tau_2$ required for completing translocation, as shown in Supplementary Figure S2. Each of the three times varies from one molecule to another; the variation is described by probability distributions. The probability distribution for the total time is the convolution of the three probability distributions for the three of its parts. Since the result of convolution is independent of the order of the parts, we can change the model by assuming that the unfolding of mDHFR is the first of the three parts, and it is followed by the translocation time $\tau_1+\tau_2$. This approach simplifies the model equations, but it has no effect on the result.

To calculate luminescence intensity, we define the functions $X_1(t)$, $Y_1(t)$, $Z_1(t)$, $X_2(t)$, $Y_2(t)$, and $Z_2(t)$, where the subscripts 1 and 2 refer to substrate proteins that were oxidized and reduced, respectively, at $t = T_0$. Functions $X_1(t)$ and $X_2(t)$ describe the molar fraction of pOA-DHFR molecules that are waiting for mDHFR to unfold and are still translocation competent at time $t$. Functions $Y_1(t)$ and $Y_2(t)$ describe the molar fraction of pOA-DHFR that have resumed translocation after mDHFR unfolding. Functions $Z_1(t)$ and $Z_2(t)$ describe the molar fraction of pOA-DHFR that have completed translocation.

Differential equations for $X_1(t)$ and $Y_1(t)$ and their solutions are

$$\begin{aligned}
\frac{dX_1(t)}{dt} &= 0 && -\infty<t<T_0 \\
\frac{dX_1(t)}{dt} &= -\alpha_2 X_1(t) && T_0\le t<T_R \\
\frac{dX_1(t)}{dt} &= -\alpha_1 X_1(t)-\alpha_2 X_1(t) && T_R\le t<+\infty
\end{aligned} \qquad \text{Eq. S2}$$

$$\frac{dY_1(t)}{dt} = 0 \qquad -\infty < t < T_R$$
$$\frac{dY_1(t)}{dt} = +\alpha_1 X_1(t) \qquad T_R \le t < +\infty \qquad \text{Eq. S3}$$

$$X_1(t) = \alpha_3 \qquad -\infty < t < T_0$$
$$X_1(t) = \alpha_3 \exp[-\alpha_2(t-T_0)] \qquad T_0 \le t < T_R \qquad \text{Eq. S4}$$
$$X_1(t) = \alpha_3 \exp[-\alpha_2(T_R-T_0) - (\alpha_1+\alpha_2)(t-T_R)] \qquad T_R \le t < +\infty$$

$$Y_1(t) = 0 \qquad -\infty < t < T_R$$
$$Y_1(t) = \frac{\alpha_3\alpha_1}{\alpha_1+\alpha_2}\exp[-\alpha_2(T_R-T_0)]\{1-\exp[-(\alpha_1+\alpha_2)(t-T_R)]\} \qquad T_R \le t < +\infty \qquad \text{Eq. S5}$$

Differential equations for $X_2(t)$ and $Y_2(t)$ and their solutions are

$$\frac{dX_2(t)}{dt} = 0 \qquad -\infty < t < T_0$$
$$\frac{dX_2(t)}{dt} = -\alpha_1 X_2(t) - \alpha_2 X_2(t) \qquad T_0 \le t < +\infty \qquad \text{Eq. S6}$$

$$\frac{dY_2(t)}{dt} = 0 \qquad -\infty < t < T_0$$
$$\frac{dY_2(t)}{dt} = +\alpha_1 X_2(t) \qquad T_0 \le t < +\infty \qquad \text{Eq. S7}$$

$$X_2(t) = (1-\alpha_3) \qquad -\infty < t < T_0$$
$$X_2(t) = (1-\alpha_3)\exp[-(\alpha_1+\alpha_2)(t-T_0)] \qquad T_0 \le t < +\infty \qquad \text{Eq. S8}$$

$$Y_2(t) = 0 \qquad -\infty < t < T_0$$
$$Y_2(t) = \frac{(1-\alpha_3)\alpha_1}{\alpha_1+\alpha_2}\{1-\exp[-(\alpha_1+\alpha_2)(t-T_0)]\} \qquad T_0 \le t < +\infty \qquad \text{Eq. S9}$$

$Z_1(t)$ and $Z_2(t)$ are calculated as convolutions of $Y_1(t)$ and $Y_2(t)$ with corresponding translocation time distributions $P_n(\tau)$, where $\tau$ is the translocation time ($\tau$ equals $\tau_1 + \tau_2$, shown in Supplementary Figure S2):

$$Z_1(t) = \int_{-\infty}^{+\infty} Y_1(t-\tau) P_1(\tau)\, d\tau \qquad \text{Eq. S10}$$

$$Z_2(t) = \int_{-\infty}^{+\infty} Y_2(t-\tau) P_2(\tau)\, d\tau \qquad \text{Eq. S11}$$

Parameterization using Gaussian distributions for $P_N(\tau)$ gives:

$$P_1(\tau) = \frac{1}{\sqrt{2\pi\alpha_5^2}} \exp\left(-\frac{(\tau-\alpha_4)^2}{2\,\alpha_5^2}\right) \qquad \text{Eq. S12}$$

$$P_2(\tau) = \frac{1}{\sqrt{2\pi\alpha_7^2}} \exp\left(-\frac{(\tau-\alpha_6)^2}{2\,\alpha_7^2}\right) \qquad \text{Eq. S13}$$

As described above, $\alpha_4$ is the mean translocation time for oxidized pOA-DHFR. $\alpha_6$ is the mean translocation time for reduced pOA-DHFR minus the time between mixing components and starting the luminescence measurement. $\alpha_5$ is the RMS deviation of translocation time for oxidized pOA-DHFR. $\alpha_7$ is the RMS deviation for the mean translocation time for reduced pOA-DHFR.

Luminescence intensity is directly proportional to $[Z_1(t)+Z_2(t)]S(t)$ where $S(t)$ is the concentration of the furimazine substrate at time $t$ divided by the concentration at $T_0$. The depletion of substrate due to enzymatic activity can be described by:

$$\frac{d\, S_{DEPL}(t)}{d\, t} = -\alpha_8 A(t) \qquad \text{Eq. S14}$$

$S_{DEPL}$ describes the time dependence of furimazine concentration, without taking into account dilutions at $t = T_G$ and $t = T_R$. $A(t)$ is the experimentally measured total luminescence intensity at time $t$. The equation above must be integrated numerically:

$$S_{DEPL}(t_n) = 1 - \frac{\alpha_8}{2} \sum_{m=1}^{n} (A_{m-1} + A_m)(t_m - t_{m-1}) \qquad \text{Eq. S15}$$

$\alpha_8$ is inversely proportional to the product of the number of furimazine molecules, quantum efficiency of the detection system, luminescence quantum yield, etc. This parameter value should be same across experiments conducted simultaneously. It was therefore linked between datasets.

The luminescence signal, $I(t)$, is obtained by subtracting background from total luminescence. $I(t)$ is directly proportional to $[Z_1(t)+Z_2(t)]S(t)$:

$$I(t) = \alpha_9 \, [Z_1(t) + Z_2(t)] \, S(t) \qquad \text{Eq. S16}$$

Here, $\alpha_9$ is a scaling factor related to instrument sensitivity that was linked between datasets.

Dilution and change in detection sensitivity between measurements (which could result from small variations in the exact positioning of the plate in the plate reader) are taken into account by $\alpha_{10}$ and $\alpha_{11}$. These parameters report the ratio of the luminescence readings after and before the first and second dilution step (at $t = T_G$ and $t = T_R$, respectively).

Furimazine concentration at each of these time intervals is thus:

$$\begin{aligned} S(t) &= S_{DEPL}(t) && -\infty < t < T_G \\ S(t) &= \alpha_{10} S_{DEPL}(t) && T_G \le t < T_R \\ S(t) &= \alpha_{11} S_{DEPL}(t) && T_R \le t < +\infty \end{aligned} \qquad \text{Eq. S17}$$

By fitting the model described above to the experimental data, we determined all values in the parameter vector $\boldsymbol{\alpha}$. Only $\alpha_8$ and $\alpha_9$ were treated as global parameters that were linked between data sets collected simultaneously. All other parameters were free to change during the fitting.

Supplementary Figure S2 provides a visual summary of the translocation model developed here. The parameter names in Supplementary Figure S2 relate to the parameter names in the description provided here as follows:

- $k_{unfold} = \alpha_1$
- $k_{incap} = \alpha_2$
- $(\tau_1 + \tau_2) = \alpha_4$

A small fraction of data points collected during luminescence measurements appeared as spikes in the signal (see Supplementary Figure S3, open circles). These data points were excluded from the final fit as follows: First, all data points without exception were fit by the model function and weighted residuals were saved. The mean squared weighted residual was calculated and multiplied by a factor of ten to produce the exclusion threshold. The points with squared weighted residuals above the threshold were then excluded from the analysis and the model was fit to the remaining data points, which produced the final fit and the values of the fitting parameters. When the error statistics are Poissonian, this method has a false positive rate of about 0.00157, i.e. about one out of 639 data points gets rejected. The false negative rate depends on the origin of the sharp spikes in the data, which was unknown, but visual inspection of the fit revealed that all spikes were removed.

#### 2. Maximum Likelihood Estimation of Cossio-Hummer-Szabo Model Parameters

The force dependence of protein unfolding rates can be extracted from the distribution of unfolding forces obtained in force-ramp experiments. Here, we describe the approach that we used to analyze the unfolding force distributions of mDHFR in the presence of ligands. The approach described here was utilized to calculate the force dependence of the folded state lifetime and its standard deviation, shown in Figure 4.

The rate of unfolding, $k$, for a folded protein in optical tweezers experiments is a function of the pulling force, $F$. How $k$ varies with $F$ depends properties of the molecule and the direction of applied force. The dependence of $k$ on $F$ and on the properties of the molecule was modeled using the formula developed by Cossio, Hummer and Szabo (Cossio *et al.*, Biophys J 111:832, 2016):

$$k(F,\boldsymbol{\alpha}) = k_0\left(1-\frac{\nu F x^{\ddagger}}{\Delta G^{\ddagger}}\right)^{2-1/\nu}\exp\left\{\frac{\Delta G^{\ddagger}}{k_B T}\left[1-\left(1-\frac{\nu F x^{\ddagger}}{\Delta G^{\ddagger}}\right)^{1/\nu}\right]\right\} \quad \text{Eq. S18}$$

Here, $\boldsymbol{\alpha} = (k_0, x^{\ddagger}, \Delta G^{\ddagger}, \nu)$ represents the vector of model parameters. $k_0$ is the unfolding rate at zero force, $x^{\ddagger}$ is the distance from the native state energy minimum to the transition state, $\Delta G^{\ddagger}$ is the barrier height, and $\nu$ is a parameter that defines the shape of the free energy potential. Values of $\nu = 1/2$ or $\nu = 2/3$ are typically used. We obtained similar $k(F,\boldsymbol{\alpha})$ for both values of $\nu$. However, as the fit was statistically better in the case of $\nu = 1/2$, we used this value of $\nu$ for our analyses.

The increase in pulling force $F$ during a force ramp measurement is determined by the trap velocity and the total stiffness of the system. Instead of modeling the time variation of the pulling force, we determined it directly from the experimental data by fitting the following function to the data:

$$F(t) = \left[\left(a(t-t_0)^{p+r(t-t_0)}\right)^{-q} + \left(F_{max}\right)^{-q}\right]^{-1/q} + c \quad \text{Eq. S19}$$

This formula was found to adequately fit the force increase during ramp experiments for force-extension curves with and without DNA overstretching. Here, $t_0$ is the time when the ramp starts, $a$ describes the slope of the ramp, $p$ describes the curvature of the force ramp, $r$ describes how the curvature changes, $q$ describes the transition from the elastic response to overstretching of DNA, and $F_{max}$ is the force at which overstretching is observed (~63 pN in our experiments). The parameter $c$ describes the starting force of the ramp.

From Eq. S19, we obtained the first derivative $F'(t) = dF(t)/dt$ and an inverse function $t = F^{-1}(f)$, such that $F[F^{-1}(f)] = f$. This made it possible to express the time derivative of the force as a function of the force itself:

$$\dot{F}(f) = F'[F^{-1}(f)] \quad \text{Eq. S20}$$

The distribution of rip forces was expressed in terms of the functions defined in Eq. S18 and S20

$$\rho[F,\dot{F}(f),\boldsymbol{\alpha}] = \frac{k(F)}{\dot{F}(F)} \exp\left[-\int_0^F \frac{k(f)}{\dot{F}(f)} df\right] \quad \text{Eq. S21}$$

To determine the unknown model parameters $k_0$, $x^‡$, and $\Delta G^‡$, the probability density in Eq. S21 must be fit to the experimental rip force data. In general, the fitting can be done using either the method of nonlinear least squares (NLLS) or the method of maximum likelihood (MML). The use of NLLS requires binning the rip forces, where the choice of bin widths affect the result. Too wide bins result in the loss of force resolution, i.e., information loss. Too narrow bins result in small counts of rip events per bin, which hinders accurate variance estimation, because rip counts follow a Poisson distribution. When the total rip count in all bins is under 10000, there is no acceptable bin width that would result in at least 100 counts per bin and at least 100 bins per distribution width. Also, binning force rip data assumes that $\dot{F}(f)$ is exactly the same for all pulling curves, which is not the case for experimental data. The MML does not require binning and therefore it is free from the problems described above.

MML searches for the values of the unknown model parameters $\boldsymbol{\alpha}$ that maximize $L(\boldsymbol{\alpha})$, the likelihood of the parameter vector $\boldsymbol{\alpha}$ given the observed data:

$$L(\boldsymbol{\alpha}) = \prod_{n=1}^{N} \rho[F_n,\dot{F}_n(f),\boldsymbol{\alpha}] \quad \text{Eq. S22}$$

Here $N$ is the total number of observed rip events, $F_n$ and $\dot{F}_n(f)$ are the rip force value and the force derivative function for the $n$-th rip event. The likelihood $L$ is often an extremely small or extremely large number, outside of the range of 64-bit floating point numbers. It is therefore more convenient to deal with the natural logarithm of $L$,

$$\ln[L(\boldsymbol{\alpha})] = \sum_{n=1}^{N} \ln\{\rho[F_n,\dot{F}_n(f),\boldsymbol{\alpha}]\} \quad \text{Eq. S23}$$

MML searches for the vector $\boldsymbol{\alpha}$ that maximizes $\ln[L(\boldsymbol{\alpha})]$. We used Newton's method to find the maximum. The method is briefly described below because one of the matrices involved in it also plays an important role in estimating the standard deviations for the model parameters and functions of these parameters. MML starts from an "initial guesses" vector $\boldsymbol{\alpha}^{(0)}$, and for this vector it calculates the elements of the vector **V** and the matrix **H**,

$$V_i = \frac{\partial \ln[L(\boldsymbol{\alpha})]}{\partial \alpha_i} \quad \text{Eq. S24}$$

$$H_{ij} = -\frac{\partial^2 \ln[L(\boldsymbol{\alpha})]}{\partial \alpha_i \partial \alpha_j} \quad \text{Eq. S25}$$

The elements of the vector and the matrix are calculated by plain summation of the derivatives of $\ln(\rho_n)$ over all $n$; where $\rho_n$ is obtained by substituting the rip force $F_n$ and the corresponding $\dot{F}_n(f)$ in Eq. S21. The negative sign in Eq. S25 is necessary to make matrix **H** positive definite, at least in the vicinity of the maximum of $\ln[L(\boldsymbol{\alpha})]$. The "correction" to the vector of initial guesses is obtained by solving the matrix equation

$$\boldsymbol{H}\,\delta\boldsymbol{\alpha} = \boldsymbol{V} \quad \text{Eq. S26}$$

The solution is obtained by first inverting the matrix **H**. The inverted matrix is denoted **C**.

The "correction vector" is then calculated

$$\delta\boldsymbol{\alpha} = \boldsymbol{C}\boldsymbol{V} \qquad \text{Eq. S27}$$

and added to the "initial guesses" vector to yield the first iteration result:

$$\boldsymbol{\alpha}^{(1)} = \boldsymbol{\alpha}^{(0)} + \delta\boldsymbol{\alpha} \qquad \text{Eq. S28}$$

The result of the first iteration plays the role of the "initial guesses" for the second iteration, and so on. Iterations continue until the vector **V** becomes very close to the null-vector, which indicates that the maximum of $\ln[L(\boldsymbol{\alpha})]$ has been reached. Taylor expansion of $\ln[L(\boldsymbol{\alpha})]$ in the vicinity of its maximum is

$$\ln[L(\boldsymbol{\alpha})] = \ln(L^{(max)}) - \frac{1}{2}\sum_{i=1}^{M}\sum_{j=1}^{M} H_{ij}\left(\alpha_i - \alpha_i^{(max)}\right)\left(\alpha_j - \alpha_j^{(max)}\right) + O[(\boldsymbol{\alpha} - \boldsymbol{\alpha}^{(max)})^3] \qquad \text{Eq. S29}$$

where $M$ is the number of unknown parameters and $O[(\boldsymbol{\alpha}-\boldsymbol{\alpha}^{(max)})^3]$ (Bachmann–Landau big $O$ notation) denotes the terms of the third and higher powers in $\boldsymbol{\alpha}-\boldsymbol{\alpha}^{(max)}$, which, according to the central limit theorem, lose significance as $N$ increases. When the number of rips $N$ is large, $O[(\boldsymbol{\alpha}-\boldsymbol{\alpha}^{(max)})^3]$ can be omitted, and then exponential function can be taken of both sides of Eq. S29, which yields

$$L(\boldsymbol{\alpha}) = L^{(max)} \exp\left[-\frac{1}{2}\sum_{i=1}^{M}\sum_{j=1}^{M} H_{ij}\left(\alpha_i - \alpha_i^{(max)}\right)\left(\alpha_j - \alpha_j^{(max)}\right)\right] \qquad \text{Eq. S30}$$

Note, that this is a general $M$-dimensional Gaussian distribution, from which it directly follows that the inverted matrix **H**, which above was denoted **C**, is the variance-covariance matrix for the estimates of the unknown model parameters. This means that the matrix **H** plays in MML exactly the same role that is played by the Hessian matrix in NLLS. The matrix **C** that was obtained as a byproduct from the last iteration makes it trivial to estimate the standard deviations of the model parameters and any functions of these parameters. The standard deviation for any model parameter equals the square root of the corresponding diagonal element of **C**,

$$\sigma\alpha_i = \sqrt{C_{ii}} \qquad \text{Eq. S31}$$

Now consider an arbitrary function Ψ (with continuous first derivatives) of the independent variable $F$ and the model parameters $\boldsymbol{\alpha}$. The standard deviation for this function can be found by linearizing it in the vicinity of $\boldsymbol{\alpha}^{(\mathrm{max})}$,

$$\sigma\Psi(F,\boldsymbol{\alpha}) = \sqrt{\sum_{i=1}^{M}\sum_{j=1}^{M} C_{ij}\left(\frac{\partial\Psi(F,\boldsymbol{\alpha})}{\partial\alpha_i}\right)\left(\frac{\partial\Psi(F,\boldsymbol{\alpha})}{\partial\alpha_j}\right)} \qquad \text{Eq. S32}$$

This approach was used to calculate the standard deviation for $\ln[\tau(F,\boldsymbol{\alpha})]$. Note, that $\ln[\tau(F,\boldsymbol{\alpha})]=-\ln[k(F,\boldsymbol{\alpha})]$, and $k(F,\boldsymbol{\alpha})$ is defined in equation S18.
